# Personalized Therapy Design for Systemic *Lupus Erythematosus* Based on the Analysis of Protein-Protein Interaction Networks

**DOI:** 10.1101/559740

**Authors:** Elizabeth J. Brant, Edward A. Rietman, Giannoula Lakka Klement, Marco Cavaglia, Jack A. Tuszynski

## Abstract

We analyzed protein expression data for Lupus patients, which have been obtained from publicly available databases. A combination of systems biology and statistical thermodynamics approaches was used to extract topological properties of the associated protein-protein interaction networks for each of the 291 patients whose samples were used to provide the molecular data. We have concluded that among the many proteins that appear to play critical roles in this pathology, most of them are either ribosomal proteins, ubiquitination pathway proteins or heat shock proteins. We propose some of the proteins identified in this study to be considered for drug targeting.

## Introduction

Systemic *lupus erythematosus* (SLE) is a unique autoimmune disease with multiple pathologies including organ damage to kidney, skin, lungs, brain and heart, among others. Women of childbearing age and African-American persons are largely affected, with a ratio of 9:1 compared to general population. Its pathogenesis includes genetics [1-4], environmental and female sex hormone [5], and epigenetics [6]. These “causal” factors result in T-, B-cell and dendritic cell dysfunctions of various types [7].

One of the important contributors to T cell dysfunction is the mitochondrial hyperpolarization, which results in ATP depletion, oxidative stress, and Ca^+2^ and actin cytoskeleton depletion [8]. The T cells eventually rupture releasing pro-inflammatory nuclear materials [8]. This causes an imbalance in the T/B cell ratio, which promotes antibody production in the peripheral blood. This whole process cascades producing excessive necrotic debris. Dendritic cells sense the excess necrotic debris and soon inflammation is out of control. A typical therapy is to target B cells to allow normal processes to remove the cellular debris [5]. A more detailed description of this pathway is given by [2, 8, 9, 10]. Here, we focus on using molecular thermodynamics methodology in conjunction with systems biology to identify key proteins in this SLE inflammatory process. This may prove to be valuable for designing new treatments on a personalized basis for this pathology and other types of diseases as has already been attempted for cancer [11-13].

### Theoretical Background

The theoretical underpinnings for the thermodynamic approach to understand the molecular biology of human diseases were developed over a several-year period and involved different examples including several types of cancer [12-17]. Here, we give a brief summary of this body of work. The transcriptome and other -omic (e.g., proteomic, genomic, etc.) measures can be viewed as representing the energetic state of a cell. By the use of the word “energetic” we mean from a thermodynamics perspective. A living system is out of thermodynamic equilibrium simply because of a constant need for metabolic energy production. It uses nutrients such as glucose and transforms them into ATP as the universal biological energy currency required for structure formation and biological function. One of the energetic demands of every cell is the production of specific proteins, which are used for numerous structural and functional need of a cell. Protein expression levels, therefore, represent a measure of the living cell’s non-equilibrium energy level. Moreover, proteins interact with other proteins generating very complex protein-protein interaction networks whose architecture is cell specific. There is a chemical potential between interacting molecules in a cell, and the chemical potential of all the proteins that interact with each other can be imagined to form a rugged landscape, not dissimilar to Waddington’s epigenetic landscape [18-19]. The above formulates our conceptual framework for the foregoing analysis.

The method we propose uses mRNA transcriptome data or RNA-seq data as a surrogate for protein concentration. This assumption is largely valid. In fact, refs. [20-21] have shown an 83% correlation between mass spectrometry proteomic information and transcriptomic information for multiple tissue types. Further, ref. [22] found a Spearman correlation of 0.8 in comparing RNAseq and mRNA transcriptome from TCGA human cancer data (https://cancergenome.nih.gov/).

Given the set of transcriptome data, a representative of protein concentration, we overlay that on the human protein-protein interaction (PPI) network from BioGrid (https://thebiogrid.org/). This means we assign to each protein on the network, the transcriptome value (or RNAseq value) after rescaling. From that we then compute the Gibbs free energy of each PPI using the standard statistical thermodynamic relationship:

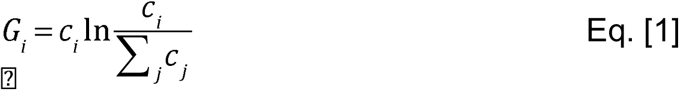

where *c*_*i*_ is the “concentration” of the protein *i*, normalized, or rescaled, to be between 0 and 1 corresponding to minimum and maximum values, respectively. The sum in the denominator is taken over all protein neighbors of *i,* and including *i.* Therefore, the denominator can be considered akin to degree-entropy as pointed out elsewhere [12-17]. Carrying out this mathematical operation essentially transforms the “concentration” value assigned to each protein to a Gibbs free energy contribution. Thus, we replace the scalar value of transcriptome to a scalar function – the Gibbs free energy.

Due to the presence of a logarithm function of a fractional number, the Gibbs free energy is a negative number, so associated with each protein on the network is a negative free energy well (local energy minimum), which corresponds to a local stability area with respect to small changes in protein concentrations. This results in a rugged free energy landscape represented schematically in Figure 1. If we use what is called a topological filtration on this landscape, we essentially move a filtration plane up from the deepest energy well. As the filtration plane is moved up, larger-and-larger energetic subnetworks are captured. For convenience, we stop the filtration at energy threshold 32 – meaning 32 nodes in the energetic subnetwork are retained. We call these subnetworks Gibbs-homology networks.

**Figure 1.**
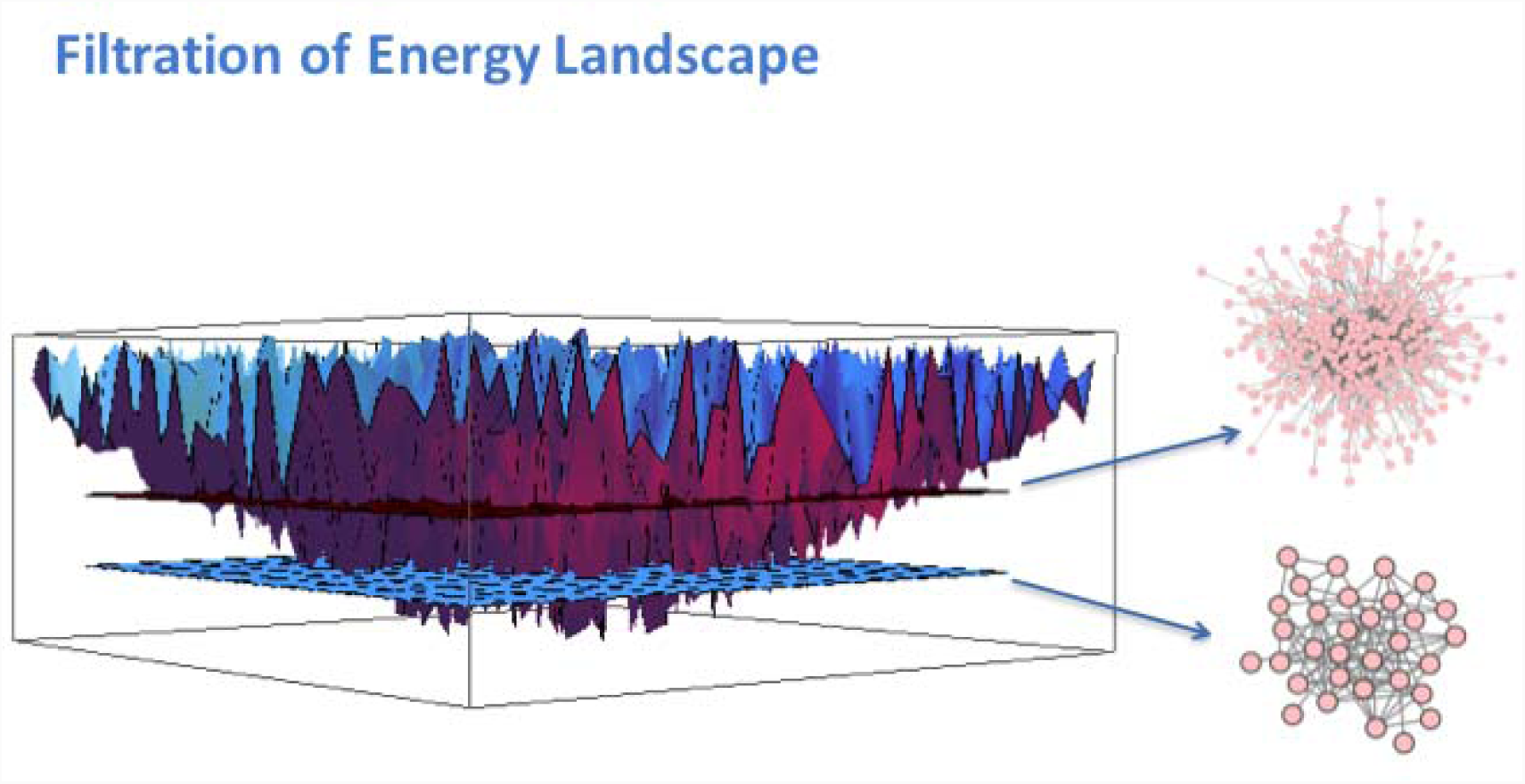
As the “filtration plane” moves up from the bottom, more-and-more nodes are captured in larger-and-larger energetic subnetworks.

We now compute the Betti centrality, which is a topological measure, of the 32-node energetic networks as described in detail earlier in ref. [15]. The main concept is easily described as follows. In networks such as PPI networks, there are holes, or rings, of various sizes. In these energetic pathways within PPI networks, the proteins form interaction rings. In densely connected, but not fully connected, networks the rings, or holes, may consist of triangles and larger rings of interaction. To find the Betti centrality we ask ourselves the following questions: which protein when removed from the network will change the overall total number of rings the most? The total number of rings is called the Betti number and is denoted *B*. Given a network *G* consisting of edges *e* and vertices *v*, the Betti centrality is given by the simple formula:

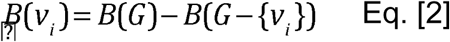

So the difference from the total Betti number *B(G)* and the Betti number of the network after removing node *i*, gives the Betti centrality for node *i*. We then compute this property for all nodes in the threshold-32 energetic network. Often there will be two or more proteins in the network that have equivalent Betti centrality making them equally important to the network. We discuss this equivalence and the Betti centrality with respect to the patient data later in this manuscript.

### Data Source and Methods

We use the Betti centrality measure, described above, on the Gibbs-homology network. The algorithm used for the calculation of Gibbs energy, Gibbs-homology and Betti centrality has been briefly described above and in detail elsewhere [13,17]. The dataset for this SLE study is from ref. [23], a study of lymphotoxin-Light pathway regulation by treatment of SLE and rheumatoid arthritis (RA) patients with baminercept. The dataset is available at GEO (https://www.ncbi.nlm.nih.gov/geo/) with the accession number GSE45219.

## Results and Discussion

As described above, we use the Gibbs-homology pathways, or small energetic networks, to find which protein is contributing most to the energetic pathway complexity. We do this by calculating the Betti number centrality. For some patients there will be one or more equivalent Betti number centralities. A Pareto chart of these “centrality proteins” for the 291 SLE patients in the GSE45291 dataset is shown in Figure 2.

**Figure 2.**
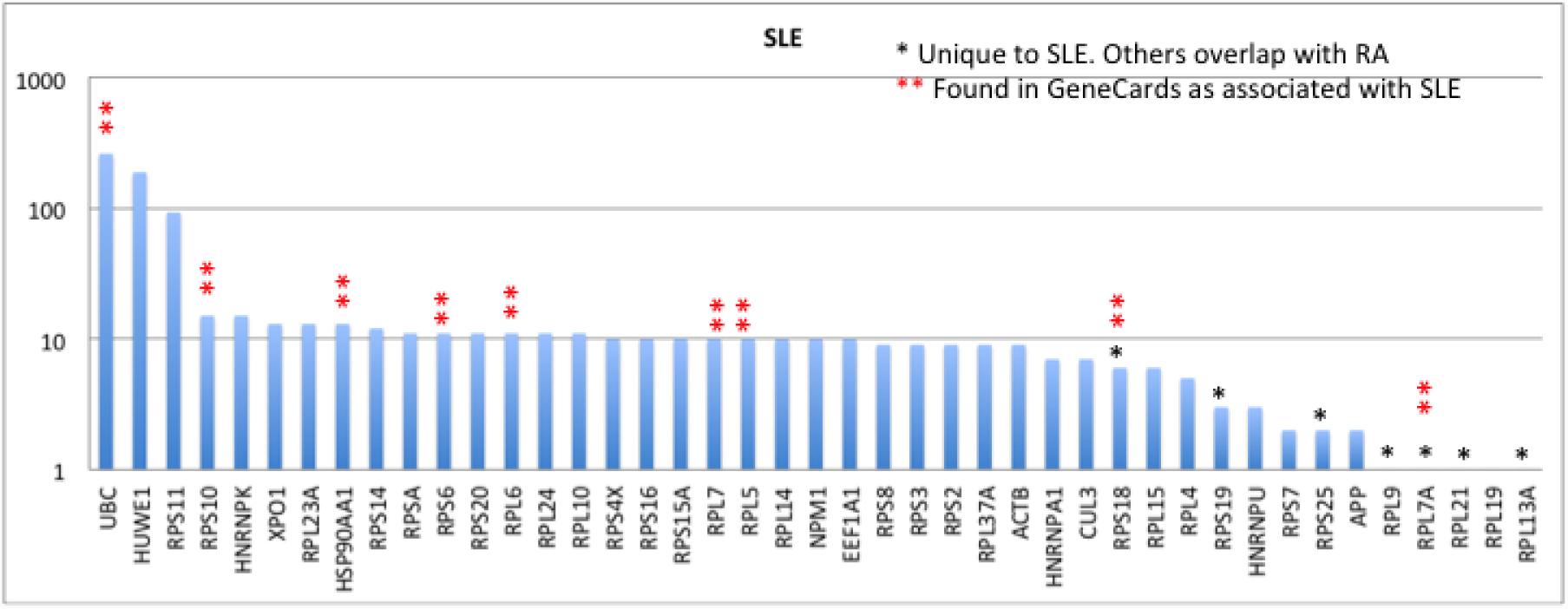
Pareto chart of highest Betti centralities for the Gibbs-homology energetic networks at threshold 32. In many networks there were one or more equivalent Betti centralities, so the total number of proteins is greater than 291, which is the number of SLE patients in the dataset. Note, the vertical axis is on a log scale.

The first thing to notice in this Pareto chart is that the number of proteins totals beyond 291 because of equivalent centralities. The second feature is that UBC, HUWE1 and RPS11 are present as centrality proteins in 100 or more patients, which is a very substantial number. Notice also the vertical axis is on a logarithmic scale, hence the actual differences are much larger than they appear.

A search of GeneCards database (http://www.genecards.org/) shows there are 1474 proteins associated with SLE. Comparing this list with a list of all the proteins in the 291 Gibbs-homology networks captured, we find an overlap of 12 proteins, namely: UBC, RPS6, RPS18, RPS10, RPLP2, RPL7, RPL6, RPL5, ISG15, IFS16, HSPA8, HSP90AA1. Interestingly and possibly importantly, these are all ribosomal proteins or proteins related to ribosomal proteins. It is worth noting that efficient ribosome biogenesis consumes over 60% of cellular energy supply in the form of ATP molecules and thus is strongly related to the energy status of the cell. This particular feature causes the nucleolar process to be highly sensitive to nutrient deprivation as demonstrated in recent studies on the target of rapamycin (TOR) signaling pathway, which plays a central role in linking the cellular nutrient status to ribosomal biogenesis [35]. In the following paragraphs we discuss each of these proteins and their role in SLE.

UBC (ubiquitin C), as to be expected, has a high Betti centrality in this population of patients, because it has high entropy. Entropy alone does not dictate high Betti centrality, but high entropy does play a role in the Gibbs energy calculation. As stated above, degree-entropy is essentially the denominator in Equation (1). The degree-entropy for UBC in the Human Biogrid protein-protein interaction network version 3.4.139 (https://thebiogrid.org/) is 1432. This means it has 1432 protein neighbors with which it interacts. Here, entropy is specifically defined as the degree-entropy or the number of interactions. Similarly, HUWE1 has 455 neighbors and RPS11 (ribosomal protein S11) has 198 neighbors. It should also be noted that UBC, HUWE1 and RPS11 are all in the ubiquitination pathway. Figure 3, shows an example of one of the networks at energy threshold 32, in which UBC is the highest Betti centrality node. In this graph the proteins in the outer ring (HUWE1, RPL10, RPS20, HNRNPU, RPS11, NPM1, RPS3, RPL5, RPS8, HNRNPK) are all neighbors. As clearly seen in this graph, UBC does not have the highest degree entropy (but RPS8 does). Nonetheless, it has the highest Betti centrality for this patient.

**Figure 3.**
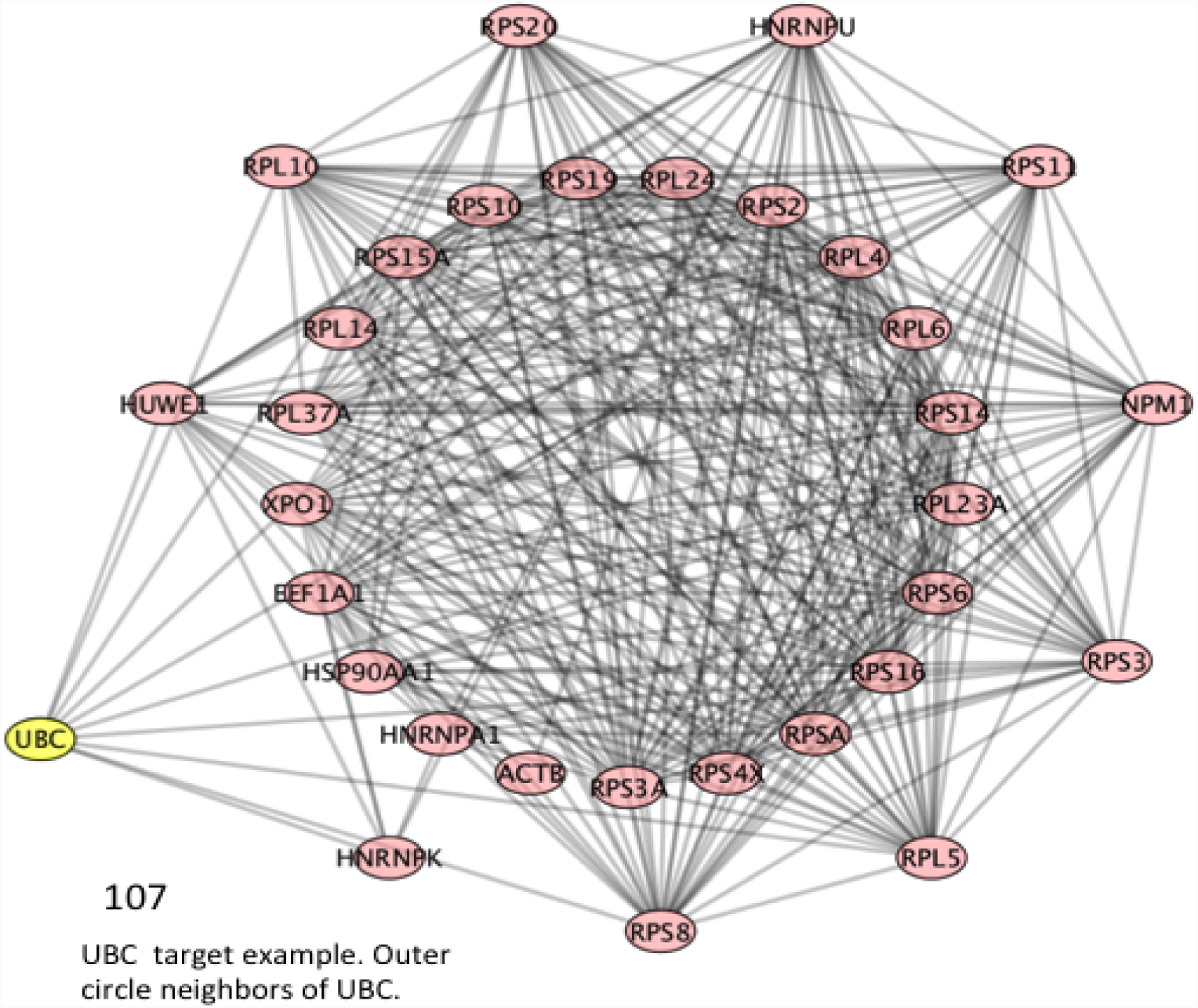
PPI Network for Patient 107, which is an example of UBC having the highest Betti centrality. The nodes in the outer ring have direct connections to UBC. The nodes in the inner ring are secondary. RPS8 has the highest number of connections.

RPS6 (ribosomal protein S6) was found in the GeneCards SLE list, and also found as having a high Betti centrality for 11 SLE patients out of the 291 total (Figure 3). However, its importance to SLE is indicated by the fact that it was found in 277 patients’ Gibbs-homology pathways at energy threshold 32. So it does not necessarily have the highest Betti centrality but it is in the “energy neighborhood” at threshold 32 for 95% of the patients. RPS6 can be phosphorylated and is associated with the functioning of mTOR in T-cell development [24]. It also plays an important role in treatment of SLE with N-acetylcysteine, a target of which is mTOR [25]. Figure 4 shows a situation where RPS6 has a high Betti centrality and in fact has three other equivalent Betti centralities (RPS20, RPSA, RPL23A). Within the nodes in the circle, CUL3 has the highest number of connections. The peripheral nodes, nodes of secondary energy importance are: HUWE1 – associated with the ubiquitination pathway; HNRNPK – known as heterogeneous nuclear ribonucleoprotein K and according to KEGG (http://www.genome.jp/kegg/pathway.html) is involved in Herpes simplex infection and viral carcinogenesis; ACTB – known as actin beta is associated with Rap1 signaling pathway and Hippo signaling pathway, platelet activation, and according to OMIM (https://www.omim.org/) is associated with Baraitser-Winter syndrome; IQGAP1 – IQ motif containing GTPase activating protein – is associated with adherens junction and regulation of cytoskeleton, and is found 34 times at energy threshold 32 in our SLE patient dataset; IFI16 – interferon gamma inducible protein number 16 – found 9 times at energy threshold 32. IFI16 is often up-regulated in SLE patients and it is suggested that it plays a key role in T-cell development. A low level of mRNA IFI16 expression has been found in naïve CD8^+^ T-cells. However, CD8^+^ mature cells express a high level of IFI16 mRNA [26]. It has also been reported that IFN-induced expression may depend on race, and the variation in coding regions of the polymorphs of the gene could account for differential regulation among individuals [26]. This implies IFI16 may be a good target for treatment by inhibition.

**Figure 4.**
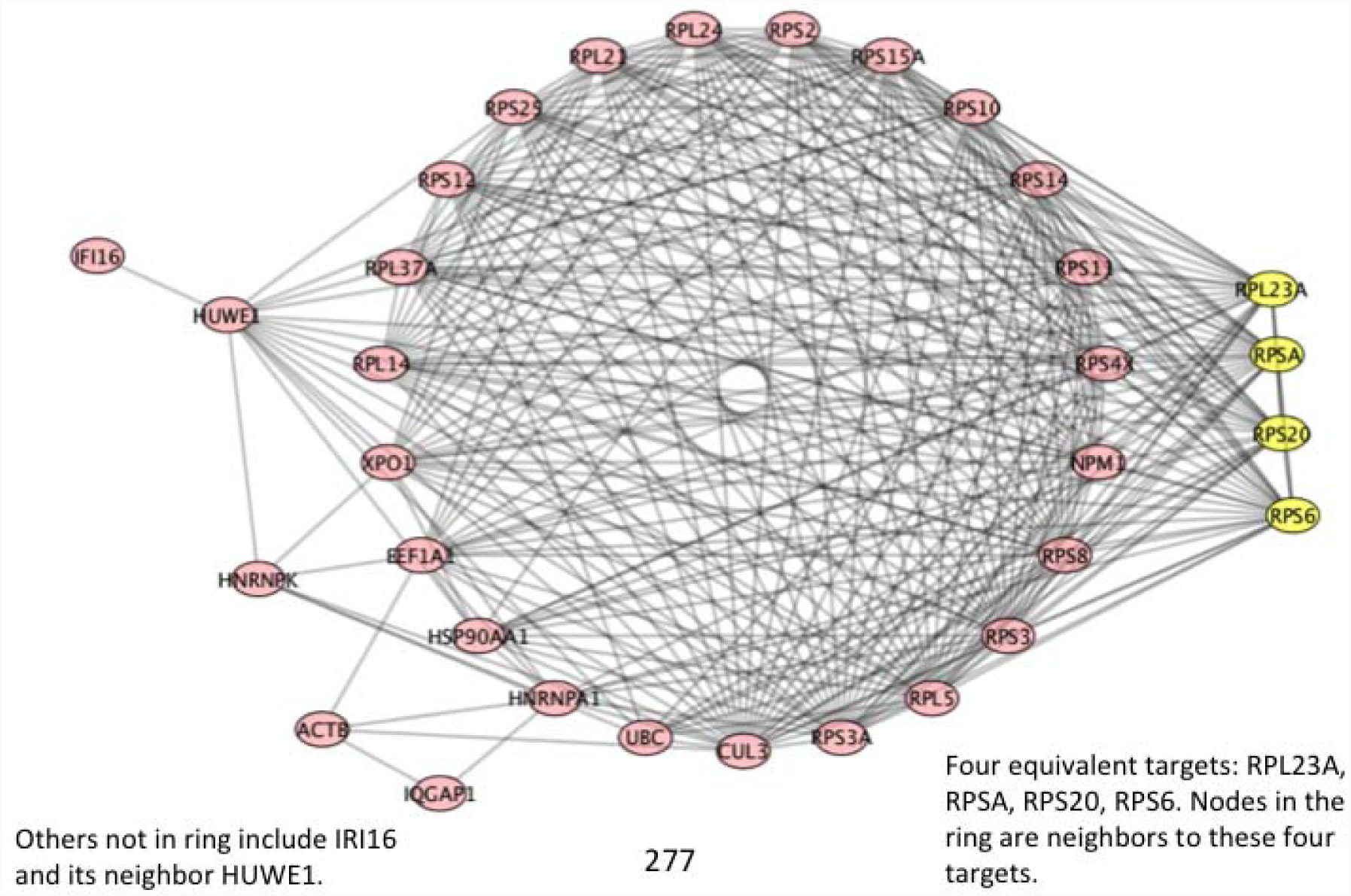
PPI Network for patient number 277. RPS6 and three other proteins (RPS20, RPSA, RPL23A) have equivalent high Betti centrality. The neighbors to these are the nodes in the circle.

Many of the proteins and high-Gibbs energy nodes discussed above are commonly found in our analysis. Consequently, we will not repeat these individual protein commentaries.

RPS10 (ribosomal protein S10) has a high Betti centrality in 15 SLE patients from our database and at Gibbs energy threshold 32 is found in 277 of the 291 patients (Figure 4). RPS10 is often dimethylated in SLE patients [27].

RPS18 (ribosomal protein S18) was found to have high Betti centrality in the Gibbs homology networks for 6 SLE patients; and at energy threshold 32, it was found in 68 networks in the 291 SLE patients (Figure 5). Incidentally, RPS6 and RPS18 are both overexpressed in cancer [28]. In this connection it is worth mentioning that ribosomal proteins are known to control the expression and activity of key tumor suppressors including p53 [36] and a mutated in multiple cancer predisposition disorders, which are known as ribosomopathies [37]. A Gibbs-homology network graph showing both RPS10 and RPS18 as equivalent high Betti centralities is shown in Figure 5. These two nodes have as nearest neighbors all the nodes in the ring. The secondary nodes are: ACTB, UBC, HNRNPA1, EEF1A1, and HSP90AA1. HNRNPA1 has a high Betti centrality 7 times in our database and is present in 278 Gibbs homology networks at threshold 32. According to KEGG it, like HNRNPK, is involved in splicesome. Further, according to OMIM (omim.org) it contributes to Paget disease and amyotrophic lateral sclerosis. EEF1A1 (elongation factor 1 alpha promoter) is very commonly associated with prostate cancer [29], among many other cancers. It is also associated with oncogenesis, apoptosis and viral infections [30]. Of course it is not surprising to see HSP90AA1 in the energetic pathway. Heat shock proteins are often over-expressed in stress situations ranging from simple lesions to cancer. Its role in lupus is discussed in ref. [31].

**Figure 5.**
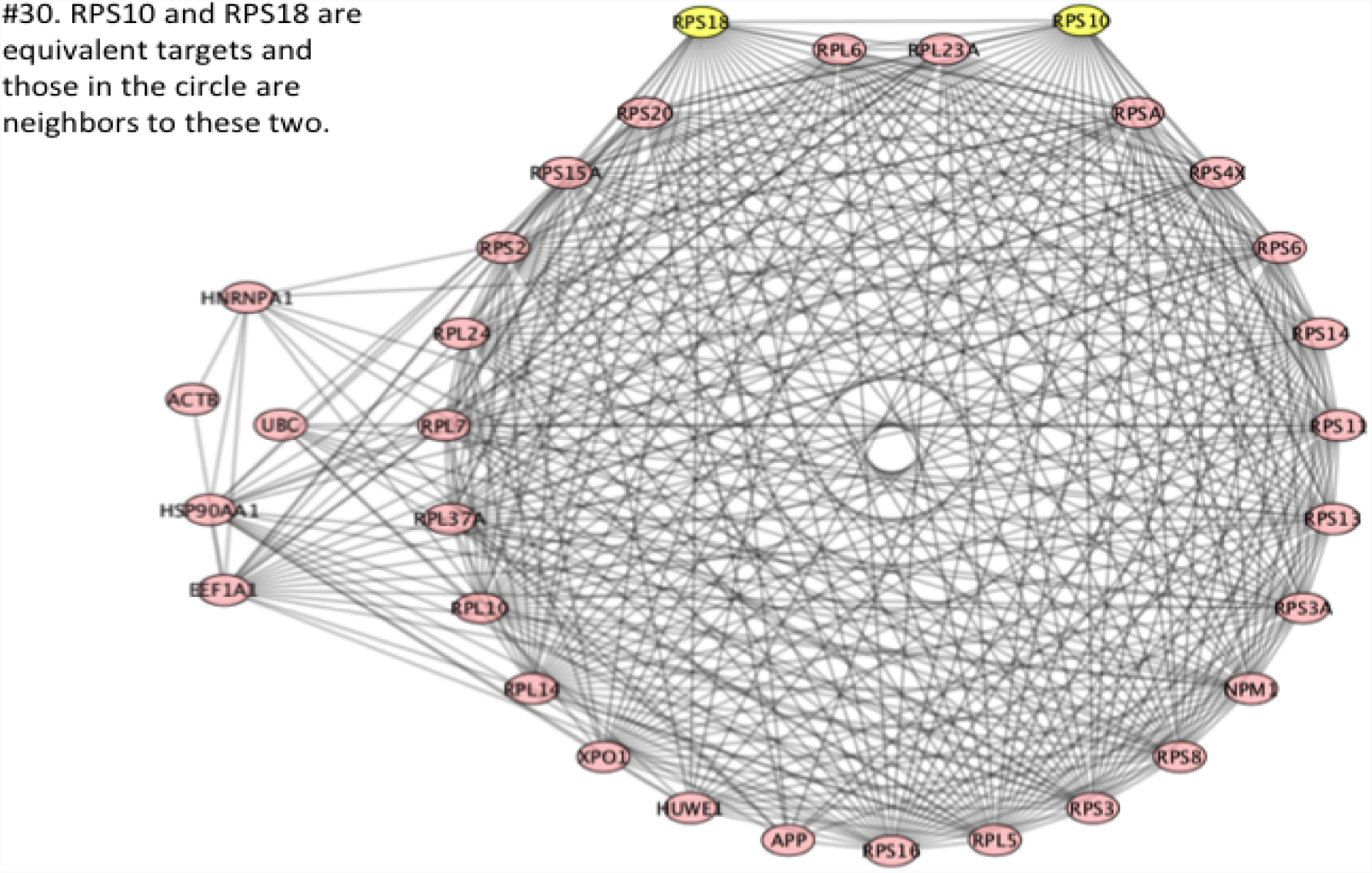
Patient number 30, RPS10 and RPS18 are equivalent high Betti centrality. The nodes in the ring are nearest neighbors to these two.

In Figure 6 we see three, high Betti centrality nodes: RPL7, RPL7A and RPS19. All the proteins in the ring are neighbors of these three. RPL7 (ribosomal protein L7) was found to have a high Betti centrality in 10 patients and found to have high Gibbs energy 252 times in the database of 291 SLE patients. It is well-known that in SLE patients an autoimmune response to RPL7 is related to T cell activity [31]. RPL7A has a high Betti centrality only once out of 291 SLE patients. Furthermore, it has a high Gibbs energy (threshold 32) in 3 patients. RPS19 has a high Betti centrality 3 times and a high Gibbs energy 37 times. Very little can be found in the literature that describes RPS19 and SLE. According to OMIM it is associated with Diamond-Blackfan anemia. The two secondary energy nodes in Figure 6 are RPS13 and RPS3A. Interestingly, neither has a high Betti centrality in the database of 291 patients. But they have a high Gibbs energy. RPS13 has a high Gibbs energy for 30 patients and RPS3A has a high Gibbs energy for 278 patients. These both are clearly important nodes for deeper investigation either as biomarkers or protein targets for inhibition.

**Figure 6.**
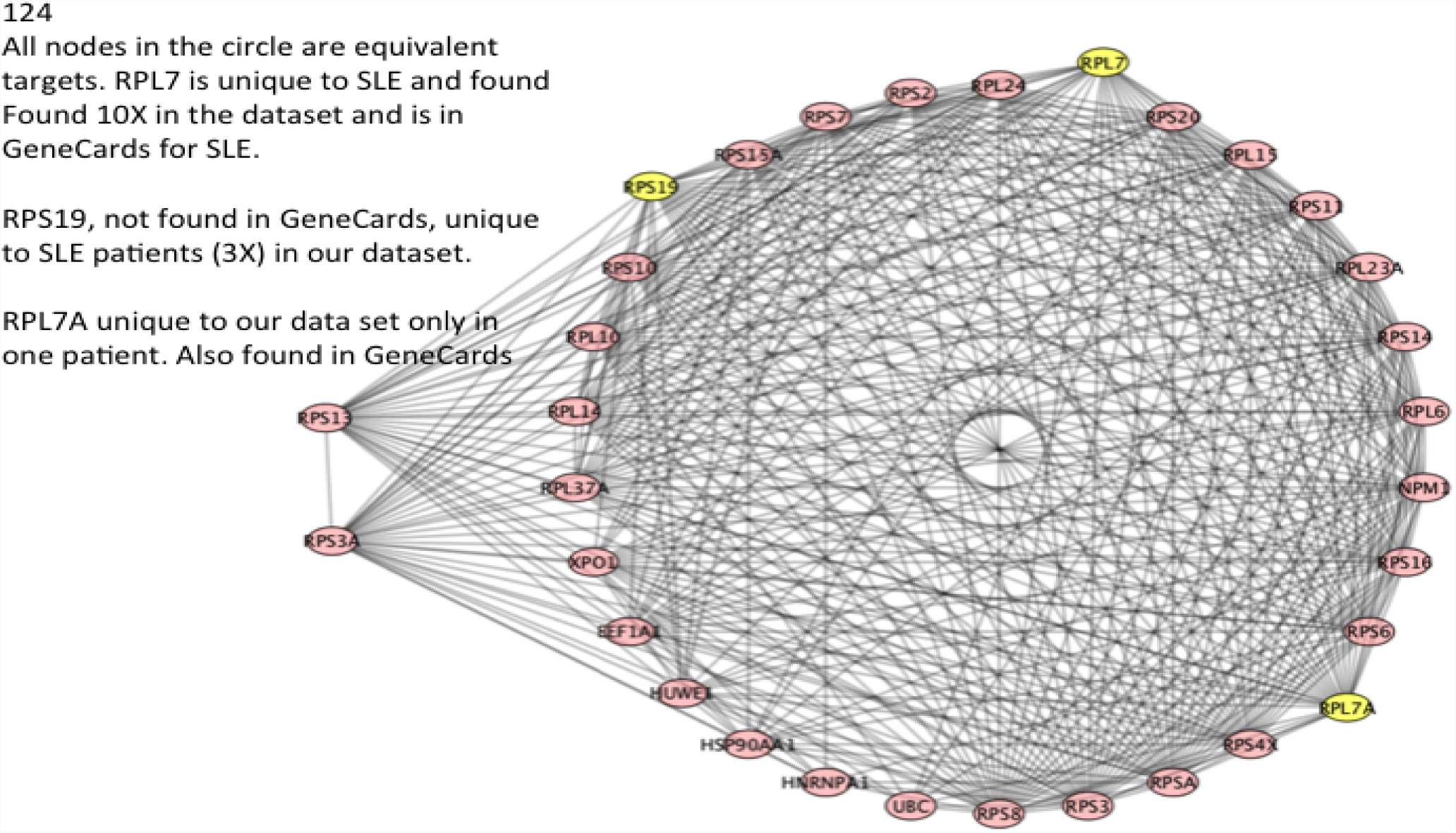
Patient number 124, all nodes in the circle have equivalent high Betti centrality. RPS19, RPL7 and RPL7A have the highest Betti centrality in this graph.

In Figure 7, RPL6 (ribosomal protein L6), and all those in the inner circle were found to have a high equivalent Betti centrality. RPL6 was found to have a high Betti centrality in 11 patients and to have a high Gibbs energy 250 times in the dataset of 291 SLE patients. It was also found in GeneCards as being important to SLE. The graph contains interferon-inducible family gene number 16 (IFI16). It is associated with SLE [26]. IFI16 was found in GeneCards as being important to SLE. We found it four times in our database of 291 patients. In patient #265 (see Figure 8) it is a neighbor of two high Betti centrality nodes, APP (highlighted) and RPL10 (not highlighted). It is also a neighbor of HUWE1 and HNRNPU. APP (Amyloid Precursor Protein) has a high Betti centrality. According to GeneCards it is not related to SLE but is associated with Alzheimer’s disease. APP is found to have a high Gibbs energy 25 times in the database of 291 patients. Lastly, we point out that IQGAP1 is also in the energetic pathway for this patient.

**Figure 7.**
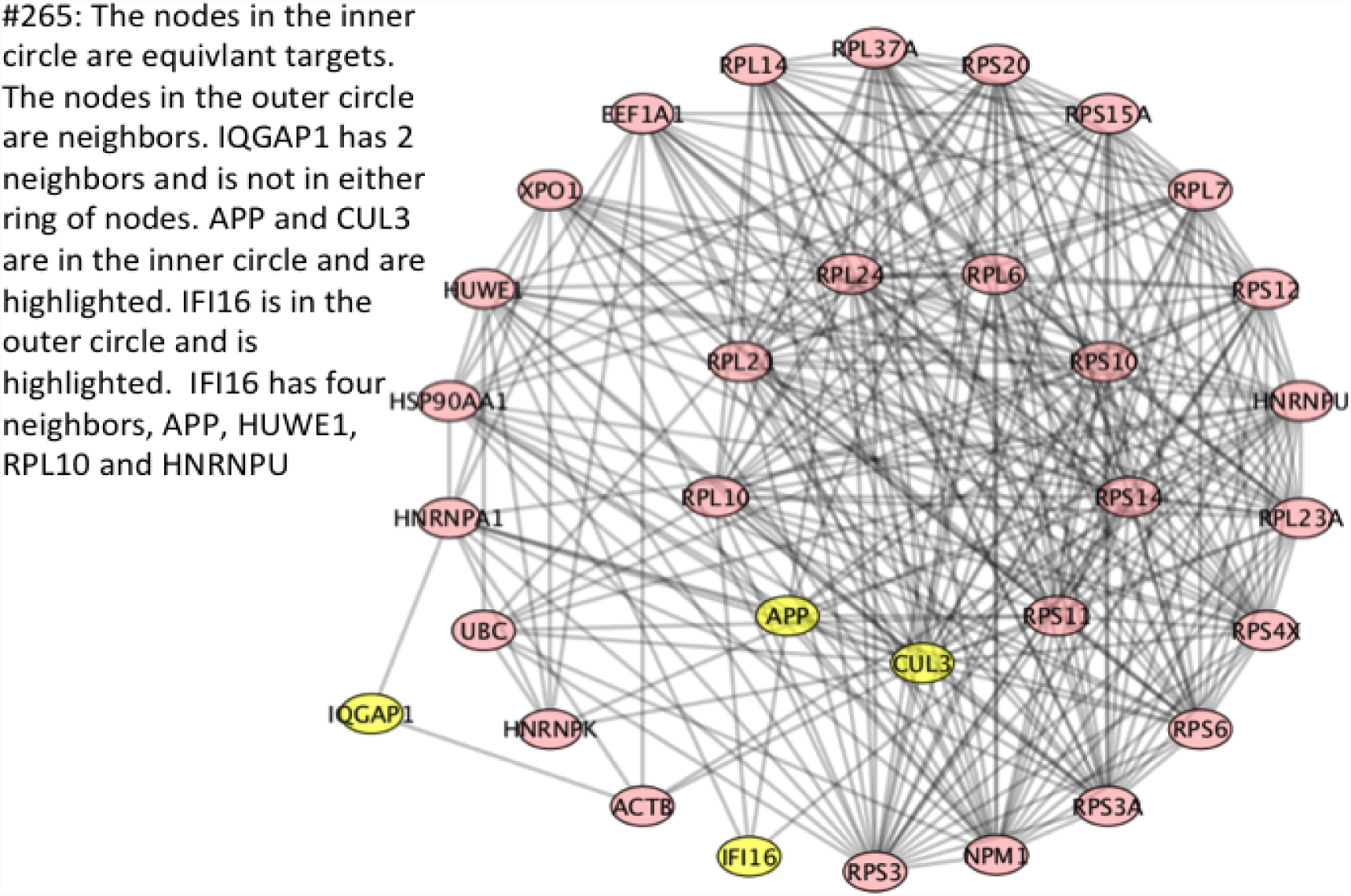
Patient number 265, RPL6, among others in the inner circle are equivalent targets.

**Figure 8.**
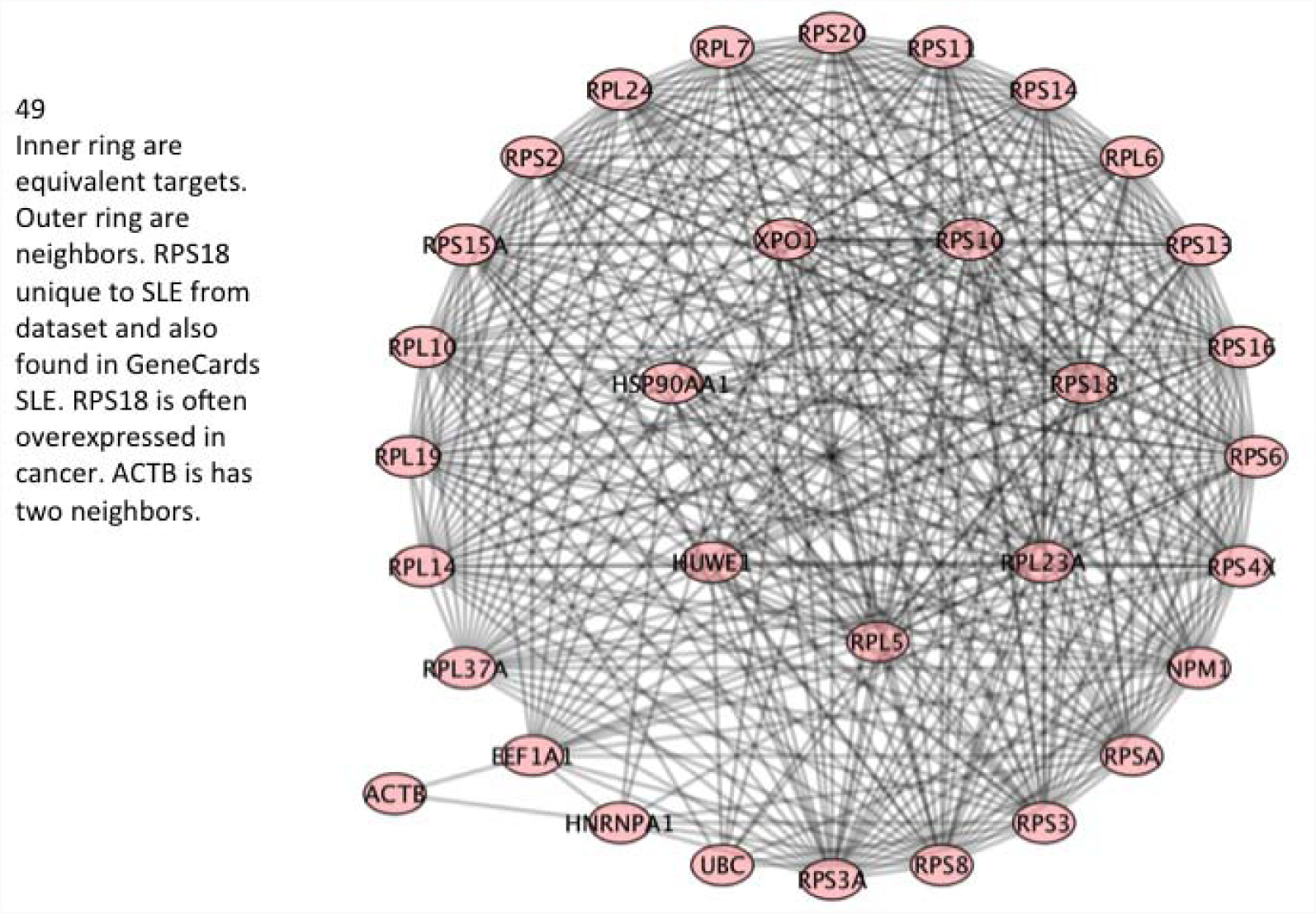
Patient 49, RPL5, among others in the inner ring are equivalent targets. Outer ring are neighbors.

RPL5 (ribosomal protein L5) was found to have a high Betti centrality in 10 patients and to have a high Gibbs energy in 21 times in the dataset of 291 SLE patients. Figure 8, patient number 49 is one of the patients with RPL5 as high Betti centrality. Very little research has been reported on RPL5, RPL6 and RPL7, with respect to SLE. Given their prevalence in our dataset (218, 250, 252 respectively), at energy threshold 32, it is reasonable to assume that these would be good proteins for further research into SLE treatment and mechanisms.

Three other proteins that should be discussed because of their frequency in our Gibbs energy analysis or because of their importance in SLE are: HSPA8, HSP90AA1 and ISG15. The two heat shock proteins HSPA8, HSP90AA1 are associated with many inflammatory diseases including rheumatoid arthritis, most cancers and SLE [32]. HSP90AA1 was found to have a high Betti centrality 13 times in the dataset of 291 SLE patients and found to have a high Gibbs energy (threshold 32) 278 times – almost every patient.

ISG15 interferon-stimulated gene 15, was found in GeneCards as being associated with SLE. More specifically, ref. [33] found it to be more highly expressed relative to healthy controls (p = 0.032) in patients with SLE who had lymphocytopenia prior to treatment. ISG15 is believed to up-regulate macrophage migration inhibitory factor [34]. Further, it has 187 protein neighbors in the version of BioGrid PPI we used. It was twice highly-expressed in our dataset of 291 SLE patients, and was found to be highly expressed in four of the 492 rheumatoid arthritis patients.

Finally, we list ribosomal proteins and their functions that are all associated with immune signaling response and were found to have high Betti centrality: RPL13A, GAIT complex formation; RPS3, activation of NFkB; RPSA, target of tuberculosis drug PZA; RPS19, inhibition of MIF, ERK and NFkB, also interacts with hantavirus; RPS6, stabilizes LANA; RPS25, promotes virus production [35]. RPL13A is a negative regulator of inflammatory proteins, thus it likely plays an important role in SLE.

## Conclusion

This paper reports the results of a computational study analyzing protein-protein interaction networks involved in Lupus, which is a unique auto-immune disease. We analyzed patient-specific data publicly available through the GEO database. We used a combination of systems biology (protein-protein interaction network analysis via Betti number calculations) and statistical thermodynamics (via Gibbs homology with energy threshold filtering) approaches to obtain information, which has statistical significance. We found close to 300 proteins which play substantial roles in the patient population of approximately the same size. However, on closer inspection, most commonly implicated proteins with major roles in the PPI networks belong to only a few special classes. The most important class consists of ribosomal proteins and ribosomal-related proteins. Next, ubiquitin and proteins belonging to the ubiquitination pathways have shown to figure prominently in this dataset. Finally, heat-shock proteins have been found to be importantly involved in these pathologies. Some of the identified proteins should be considered for therapeutic inhibition as they clearly appear to be biomarkers for Lupus.

## Acknowledgments

EAR and GLK acknowledge partial funding from CSTS-Healthcare, Toronto, Canada. JAT acknowledges funding support from NSERC (Canada).

## Author Contributions

EJB conceived of the idea and suggested it. EAR did the analysis, EJB and GLK provided clinical and biomedical insight. JAT contributed to the methodology development. MC assisted in interpreting the results. All authors contributed to writing the manuscript.

